# Catestatin regulates the colonic mucus layer in inflammatory bowel disease

**DOI:** 10.1101/2021.02.09.430377

**Authors:** Elke M. Muntjewerff, Lisanne Lutter, Kechun Tang, Mariska Kea-te Lindert, Jack Fransen, Bas Oldenburg, Sushil K. Mahata, Geert van den Bogaart

## Abstract

**Background:** The pro-hormone chromogranin A (CgA) and its bioactive cleavage product catestatin (CST) are both associated with inflammatory bowel disease (IBD) and dysregulated barrier functions, but their exact role has remained elusive. Here, we demonstrate that CST regulates the colonic mucus layer.

**Methods:** CST levels were measured in feces of IBD patients. The mucus layer, goblet cells, and immune cell infiltration were analyzed by histology and electron microscopy in colon tissue from IBD patients and mice with selective deletion of the CST-coding region of the CgA gene.

**Results:** CST levels were elevated in feces of IBD patients compared to healthy controls. The thickness of the mucus layer was increased in non-affected, but not in inflamed, regions of the colon in IBD patients. The thickness of the mucus layer and concomitant mucus production were also increased in the CST-KO mouse. This mucus phenotype in CST-KO mice could be reversed by bone marrow transplantation from wildtype mice.

**Conclusions:** CST produced by bone-marrow derived immune cells reduces production of the mucus layer in the intestine. This might contribute to the reduced mucus layer in inflamed colon regions of IBD patients. Additionally, CST feces levels might be a biomarker for IBD.

## Introduction

Approximately two million people in the United States suffer from IBD, comprising Crohn’s disease (CD) and ulcerative colitis (UC)^1^. In IBD patients the plasma levels of both the pro-hormone chromogranin A (CgA)^2–6^ and its bioactive cleavage product catestatin (CST: human CgA_352-372_)^6–8^ are increased. CST is generally considered anti-inflammatory^9,10^ and treatment of UC mouse models with CST was found to reduce macrophage infiltration of the colon and attenuate inflammation^11–13^. We recently showed that CST-KO mice, with a selective deletion of the CST-coding region of the *Chga* gene, display severely compromised intestinal permeability and inflammation in the colon^6^. Additionally, the colon of CST-KO mice displays increased fibrosis, macrophage infiltration and a bacterial population reminiscent of IBD and/or inflammatory bowel syndrome patients^6^. These inflammatory and permeability phenotypes are reversible by administration of exogenous CST^6^. However, the exact role of CST in mucus regulation has not yet been addressed.

In addition to the epithelial cellular junctions, the mucus layer contributes to the protective barrier function of the colon^14^. The mucus is produced by the goblet cells and consists of antimicrobial enzymes, immunoglobulins, glycoproteins and mucins (MUC)^14^, with the latter two components defining the mucus layer composition. MUC2 is particularly important for mucus organization in the colon, since it establishes the connection between the compact inner layer and outer, more loose, mucus layer^15^. The outer layer interacts with bacteria, while the inner layer is impenetrable and protects the mucosa from bacterial infiltration^15^. A damaged mucus layer exposes the intestinal epithelial cells to gut microbial components which might lead to colonic inflammation^14^. In UC, the mucus barrier is damaged independently of local inflammation, suggesting an intrinsic defect of the mucus layer^16^. Mucus production is generally less affected in CD, but patients still display bacterial infiltration, again indicating that the mucus composition is not sufficient to maintain (mucosal) homeostasis^14,17^.

Aim of this study was to identify how CST affects the colonic mucus layer. We characterized the thickness of the mucus layer in non- and inflamed regions of the colon of CD patients, inflamed colon of UC patients, and healthy controls. Moreover, we measured the CST concentrations in feces of IBD patients. We also characterized the thickness of the mucus layer in CST-KO and wildtype (WT) mice. Since we recently showed that macrophages (and possibly other bone-marrow derived immune cells) are major producers of CST^18,19^, we determined the effect of bone-marrow transfer from WT mice into CST-KO mice on the colonic mucus layer.

## Materials and Methods

### 1.1 Mice

Male WT, CgA-KO and CST-KO mice (3 months old) on C57BL/6J background were used to study the intestine. All procedures relating to the WT and KO mice were approved by the Institutional Animal Care and Use Committee (UCSD) and all procedures and housing and handling of the animals were in accordance with National Institutes of Health animal care guidelines. CST-KO have been described before^19^. CST-KO mice display hypertension, and a hyperadrenergic and insulin-resistant phenotype. Bone-marrow transplant from WT to CST-KO mice was performed as described previously^18^.

### 1.2 Human colon biopsies and feces collection

Colon biopsies and feces samples were collected from patients with CD (N=39) and UC (N=36) (Table S1) in a cohort of long-standing CD and UC patients stratified for inflammation by colonoscopy. A flare-up or inflammation was defined as histological and endoscopic disease activity, whereas remission entailed absence of endoscopic inflammation and of histological disease activity. Non-inflamed biopsies were taken of the same colon segment > 2 cm from endoscopically visible disease. Feces from healthy controls (N=10) was also collected (average age 44 with range 38-54 years, 7/10 female) (Table S1). Informed consent was obtained and the study was conducted in accordance with the Institutional Board Review of the University Medical Center Utrecht (approval number NL 35053.041.11 & 11-050 for the IBD cohort and NL40331.078 for the healthy controls). Samples were collected in compliance with the Declaration of Helsinki.

### 1.3 CgA, CST and calprotectin levels in human feces

Feces was used to determine CgA and CST levels using ELISA (CUSABIO CSB-E17355h, CSB-E09153h) and calprotectin (HycultBiotech HK379).

### 1.4 Electron microscopy mouse colon

Human colon was fixed with 2% glutaraldehyde and 2% paraformaldehyde in 0.1 M cacodylate (pH 7.4) buffer overnight at 4°C. After washing in buffer tissue was postfixed for 1 h at RT in 1% osmium tetroxide and 1,5% potassium ferrocyanide in 0.1 M cacodylate buffer, washed in MQ, followed by one hour staining in *en bloc* with 2% uranyl acetate on ice. After washing in MQ, tissue was dehydrated in an ascending series of aqueous ethanol solutions and subsequently transferred via a mixture of aceton and Durcupan to pure Durcupan (Sigma) as embedding medium. Ultrathin sections (80 nm) were cut on a Leica Artros ultramicrotome, picked up on grids, stained with 2% uranyl acetate and lead citrate, air dried and examined in a JEOL JEM1400 electron microscope operating at 80 kV.

### 1.5 Electron microscopy human colon

Human colon was fixed with 2% glutaraldehyde in 0.1 M cacodylate (pH 7.4) buffer overnight at 4°C. Tissue was postfixed for 1 h at RT in 1% osmium tetroxide and 1.5% potassium ferrocyanide in 0.1 M cacodylate buffer, washed in MQ, followed by one hour staining in en bloc with 2% uranyl acetate on ice. After washing in MQ, dehydrated in an ascending series of aqueous ethanol solutions and subsequently transferred via a mixture of aceton and Durcupan to pure Durcupan (Sigma) as embedding medium. Ultrathin sections (80 nm) were cut on a Leica ultramicrotome, and picked up on grids, stained with 2% uranyl acetate and lead citrate, air dried and examined in a JEOL JEM1400 electron microscope (JEOL) operating at 80 kV.

### 1.6 Processing and histology human & mouse colon

Human colon was fixed in carnoy-methanol to preserve the mucus layer. Mouse colon was fixed with 2.5% glutaraldehyde in 0.15 M cacodylate buffer. Mouse and human colon was dehydrated in ethanol series, xylene and paraffin. Afterwards, the tissue was carefully embedded in paraffin (formalin fixed and paraffin embedded (FFPE)) and sections of 5 μM were cut. Hematoxylin and eosin (HE) and periodic acid-Schiff (PAS) staining were performed using standard techniques. Slides were mounted on silane coated glass slides (MUTO PURE CHEMICALS 511618) using Quick D mounting medium and imaged using the PerkinElmer Vectra (Vectra 3.0.3; PerkinElmer, MA) at 20x magnification.

### 1.7 Analysis Electron microscopy goblet cells and mucus layer in colon

The imageJ straight line tool was used to analyze the microvilli (from border till visible end in lumen) and mucus thickness (measured on top of microvilli). Additionally, the freehand drawing tool was used for measuring the goblet cell area.

### 1.8 Statistical data analysis

Data are expressed as mean ± SEM. D’Agostino & Pearson omnibus normality test was performed followed by one-way ANOVA with Bonferroni post-hoc tests or the Kruskal-Wallis with Dunn’s. Significance is displayed in the graphs and a value of p < 0.05 was considered statistically significant.

## Results

### Feces CST correlates with IBD activity

We recently showed that CST is elevated in plasma of CD patients regardless of disease state, flare-up or remission, compared to healthy controls^6^. Here, we addressed whether CST is also elevated in the feces of IBD patients. We measured CgA and CST levels in feces samples from a cohort of long-standing CD and UC patients stratified for inflammation by colonoscopy and confirmed by calprotectin levels (Fig. S1A-B, Table S1). For CD, CgA feces levels were only increased in patients with flare-ups, whereas for both CD and UC, CST feces levels were increased regardless of disease state (Fig. 1, Table S1). These findings are reminiscent to our previous findings in plasma^6^ and support the conclusion that CST is not only elevated in circulation, but also locally in the colon of IBD patients.

**Fig. 1.**
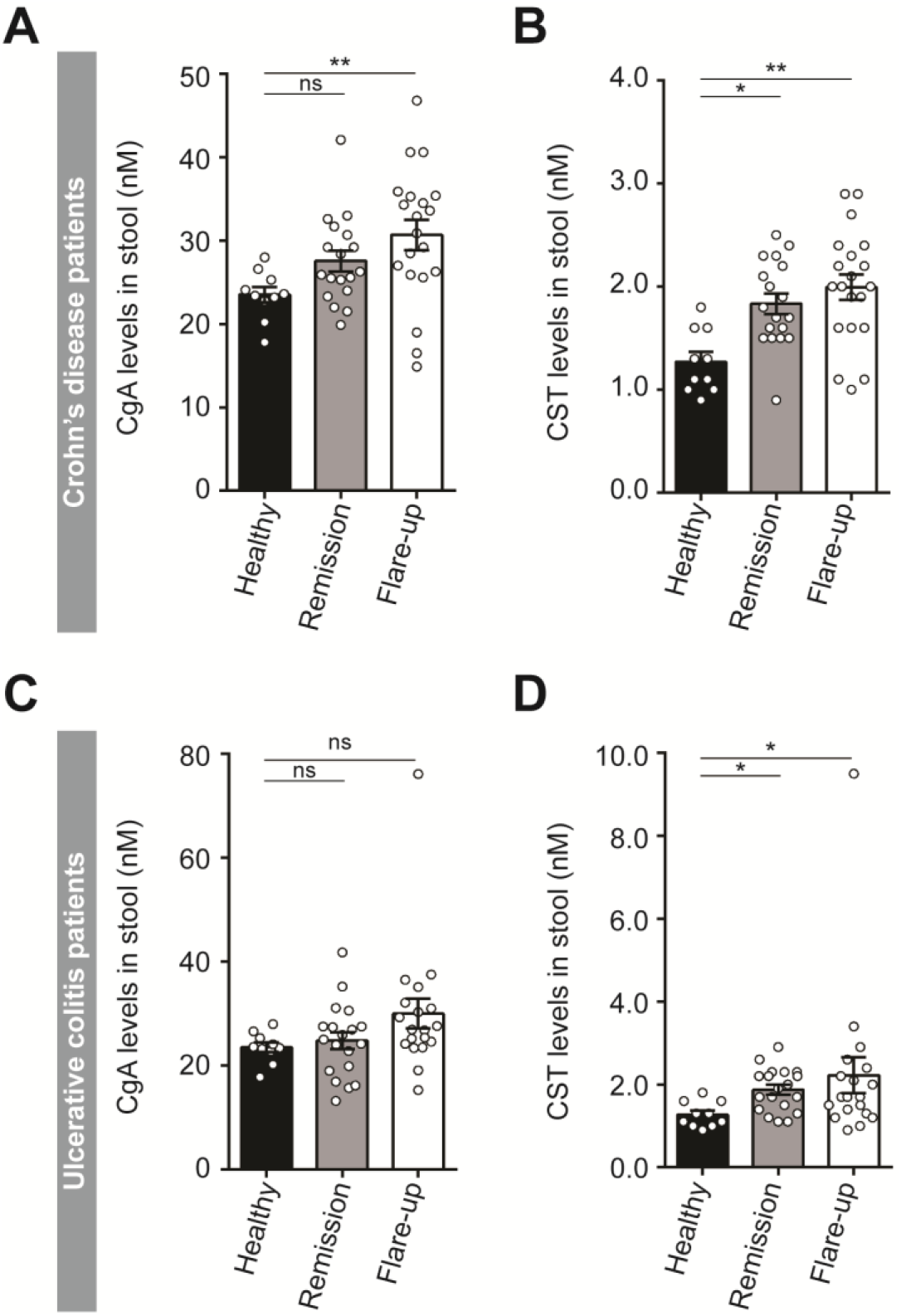
CST correlates with IBD activity. **(A)** CgA levels in feces of healthy donors (black) and Crohn’s disease (CD) patients in remission (grey) or flare-up (white). **(B)** Same as panel A, but now for CST levels in feces. **(C)** Same as panel A, but now CgA levels for Ulcerative Colitis (UC) patients. **(D)** Same as panel C, but now for CST. Kruskal-Wallis *: P<0.05; **: P<0.01; ***P<0.001.

### Mucus layer thickness depends on local inflammation

In order to address the relation between inflammation and the mucus layer, we compared non-inflamed and inflamed regions of colon biopsies of IBD patients (Fig. 2A-B, Fig. S2). Using cytochemistry and electron microscopy, we observed an increased thickness of the mucus layer at non-inflamed sites of the colon in IBD patients, while the thickness in inflamed regions was unaltered compared to a healthy individual (Fig. 2C-G). Moreover, electron microscopy revealed that the microvilli were shorter in inflamed as compared to non-inflamed regions of the colon within the same patients (Fig. 2C-F). Especially for CD patients, non-inflamed regions displayed longer microvilli and increased mucus thickness (Fig. 2C-E, Table S2). The thickness of the protective mucus layer of CD patients thus depends on the inflammatory state of the mucosa.

**Fig. 2.**
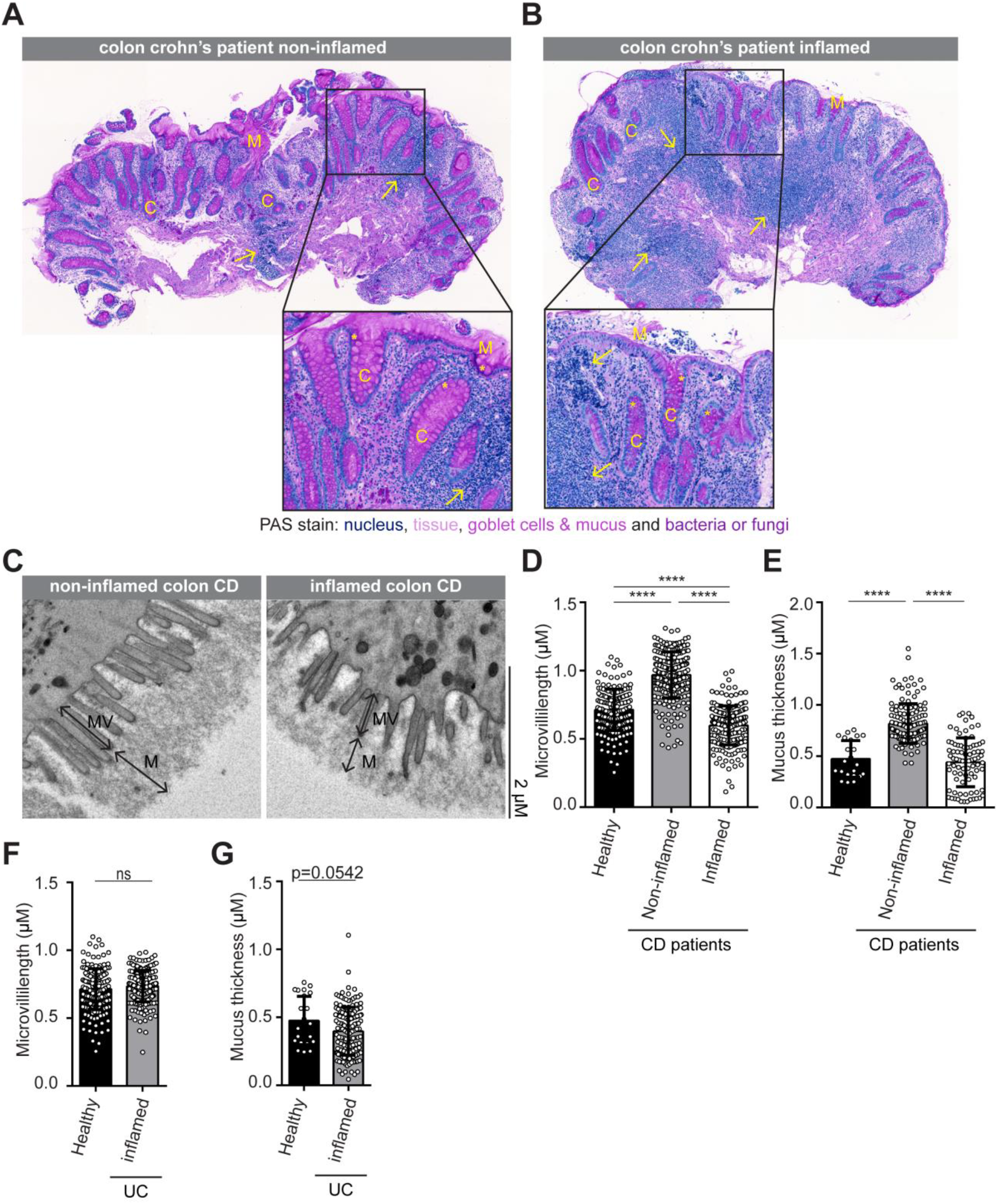
IBD inflammation links to mucus layer thickness. **(A-B)** Representative images of periodic acid–Schiff (PAS) stained colon cross-sections of non-inflamed **(A)** and inflamed **(B)** colon tissue of Crohn’s disease (CD) patients with enlargement in black framework. M: mucus, C: crypt, *: goblet cell and arrows indicate immune cell infiltration. **(C)** Representative electron microscopy (EM) micrographs of colon showing mucus layer (M), microvilli (MV) and epithelial border (EB). **(D-G)** Quantification of microvilli length and mucus thickness in colon of Crohn’s disease (CD) **(D-E)** and Ulcerative Colitis (UC) **(F-G)** patients. Kruskal-Wallis *: P<0.05; **: P<0.01; ***P<0.001; ****P<0.0001.

### CST affects goblet cells and promotes mucus production

In the next set of experiments, we addressed the question whether the altered mucus layer in IBD patients could be linked to elevated levels of CST. In the colon of 3 months old CST-KO mice, the thickness of the mucus layer and the length of the microvilli were increased when compared to WT mice (Fig. 3A-C). The number of goblet cells was also increased in the colon of CST-KO mice (Fig. 3D-E), and electron microscopy revealed an altered morphology of these goblet cells (Fig. 3F-H). Quantitative PCR showed that expression of MUC2 was increased in the CST-KO mice (Fig. 3I). This increased MUC2 expression was fully reversible by intraperitoneal injection of CST (Fig. 3I).

**Fig. 3.**
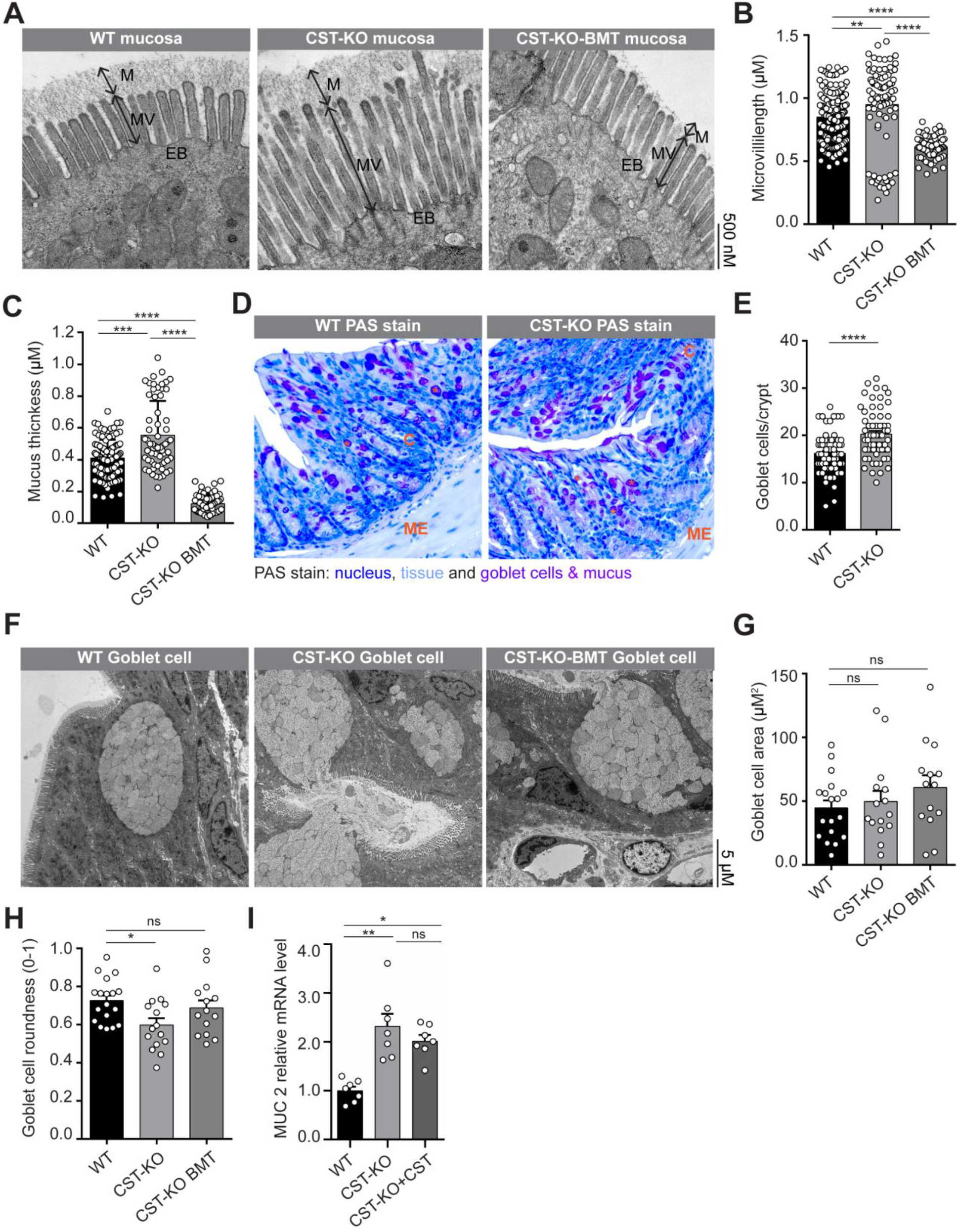
CST affects goblet cells and mucus production. **(A)** Representative electron microscopy (EM) micrographs of colon from wildtype (WT), CST-KO and CST-KO mice that received a bone marrow transplant from WT mice (CST-KO-BMT). Images show mucus layer (M), microvilli (MV) and epithelial border (EB). **(B-C)** Quantification of microvilli length **(B)** and mucus thickness **(C)** from panel A. **(D)** Representative images of periodic acid–Schiff (PAS) stained colon cross-sections of WT and CST-KO mice. ME: muscularis externa, M: mucus, C: crypt, *: goblet cell. **(E)** Quantification of goblet cells per crypt from panel D. **(F)** Electron microscopy micrographs of colon showing goblet cells. **(G-H)** Quantification of goblet cell area **(G)** and circularity **(H)** from panel F. **(I)** Relative mRNA level of *MUC2* for WT, CgA-KO, CST-KO and CST-KO treated with CST. Kruskal-Wallis *: P<0.05; **: P<0.01; ***P<0.001; ****P<0.0001.

### Bone-marrow derived immune cells produce CST in the colon

We recently showed that macrophages are major producers of CST and that macrophage-produced CST is both sufficient and required for normalization of the blood pressure in CST-KO mice^19^. CST-KO mice are hypertensive due to increased production of catecholamines, and this hypertensive phenotype could be fully reversed by depletion of macrophages or by bone-marrow transfer from WT mice to CST-KO^19^. Bone-marrow transfer in CST-KO mice was also found to restore the phenotype of the mucus layer by reducing its thickness and the length of the microvilli (Fig. 3A-C). These results indicate that CST produced by macrophages (or other bone-marrow immune cells) regulates the colonic mucosa.

## Discussion

In this study, we showed that CST is increased in the feces of IBD patients regardless of disease activity, whereas feces levels of its precursor CgA are only increased in CD patients with active disease. We also found that, compared to a healthy control, the thickness of the mucus layer and the length of the microvilli are increased in non-inflamed colon regions of CD patients, whereas the mucus layer is unaltered and the microvilli are shorter in inflamed regions. CST-KO mice display an increased thickness of the colonic mucus layer as well, in line with the increased presence and altered morphology of the goblet cells and increased levels of transcription of Muc2, a major component of the mucus layer^15^. These microvilli and mucus phenotypes could be fully reversed by bone-marrow transfer from WT cells. From these findings, three important conclusions can be drawn:

First, not only blood levels^6^, but also feces levels of CgA and CST might be markers for disease severity of IBD, particularly CD. CST levels are increased both in plasma^6^ and feces (this study) of IBD patients irrespective of disease activity, whereas levels of CgA, the precursor of CST, are only elevated during active CD. These findings suggest a proteolytic switch, where CgA and CST are excessively produced in IBD regardless of the disease state, but CgA is cleaved more into CST in quiescent disease. As we recently reported, CST inhibits colon inflammation and promotes the intestinal epithelial barrier function by counteracting pancreastatin, another cleavage product of CgA^6^. Thereby, the elevated CST levels in IBD patients are suggestive of a feedback mechanism to counteract the disease state and might help to normalize the colon function^6^.

Second, we found that the mucus layer thickness and microvilli length in IBD patients depend on local disease activity in the colon. Between 1970 and 1980, several studies showed increased goblet cell numbers in the ileum and colon^20,21^ and increased mucus secretion in the ileum of CD patients^21^. In contrast, UC patients were characterized with decreased goblet cell presence and glycocalyx (inner mucus layer)^20,22,23^. These outcomes are based on case reports and did not include non-inflamed intestine of the same patient. In our study, non-inflamed colon regions of CD patients were characterized with a thicker mucus layer than inflamed colon regions from the same patient, whereas we did not observe significant differences for UC patients. Since the patient biopsies were taken only a few centimeter apart, this indicates tight local mucosal regulation. Moreover, the mucus layer in non-affected colon regions of CD patients was even thicker than for healthy controls, indicating excessive mucus production, possibly as a compensatory mechanism to protect the colon from microbial infiltration. The length of the microvilli was affected in the same way: increased in non-inflamed tissue and shortened in inflamed colon tissue, a pattern observed before in pigs^24^. Previously, electron microscopy studies reported destroyed colon microvilli in UC patients^20,23^, whereas no consistent changes were found in CD patients^20,22^. Moreover, in the ileum of CD patients, a decreased microvilli density has been reported, as well as downregulation of genes related to microvilli formation, F-actin bundling, membrane cytoskeleton cross-linking and intermicrovillar adhesion complex^25^. Our data add to this and show that changes of the mucus layer and microvilli in the colon depend on the local inflammation state in CD patients.

Third, we conclude that CST produced by bone-marrow derived cells regulates the production of the protective mucus layer. We found that the thickness of the mucus layer in the colon of CST-KO mice is increased, in line with the increased infiltration of immune cells reported previously^66^.

It seems contrasting that CST presence reduced the protective mucus layer, but at the same time reduces intestinal permeability as we showed previously^6^. So far, we don’t know whether the composition of the decreased mucus layer in presence of CST would be more protective against bacterial infiltration. Another open question is whether the effects on mucus regulation might be caused by CST alone or by the balance between CST and pancreastatin (PST)^26^, another cleavage product of CgA. Our previous research showed that the intestinal barrier permeability is regulated by the antagonistic role of CST and PST, where CST reduced while PST increased permeability^6^. We found that the increased length of the microvilli and the increased thickness of the mucus layer in the colon of CST-KO mice were fully reversible by bone-marrow transfer from WT mice. Combined with our previous findings that macrophages produce CST themselves and that bone-marrow transfer from WT mice into CST-KO mice results in near-normal levels of CST in the plasma^6^, this suggests that the macrophages (and possibly other bone-marrow-derived cells) are major producers of CST in the colon. In line with our hypothesis that CST produced by infiltrating macrophages reduces mucus production, we found a reduced thickness of the mucus layer in inflamed regions of the colon of CD patients, which have a high density of infiltrated macrophages and other immune cells^27,28^. Further supporting this hypothesis, intraperitoneal injection of CST in CST-KO mice could reverse the expression of *MUC2*. Thus, based on the findings from this study, we expect that the elevated levels of CST produced by immune cells regulate the production of mucus in inflamed colon regions of CD patients.

An open question is what drives the cleavage of CgA to produce CST. Candidates are cathepsin L, an endolysosomal protease expressed by endocrine cells that co-localizes with CgA in chromaffin granules^29^, and plasmin, the main enzyme of the fibrinolytic cascade^30^. The expression of cathepsin L is increased in macrophages during mucosal inflammation of IBD patients^31^, as well as in the colon of CST-KO mice^6^. For plasmin, it has been shown that its inhibition prevents disease progression in acute colitis mouse models^32^. However, its expression is downregulated in CST-KO mice^6^. Finally, it seems possible that CgA is not only converted to CST by a host protease, but also by proteases produced by the gut bacteria^33^, which would mean that the micro-environment in the gut defines the balance between CgA and CST.

Overall, our finding that CST regulates the mucus layer in the gut strengthens the emerging concept^6,31,32^ that CST could be a potential new biomarker and/or therapeutic target for treatment of IBD and other gastrointestinal diseases.

## Conclusion

CST produced by bone-marrow derived immune cells reduces production of the mucus layer in the intestine. Compared to healthy controls, IBD patients have increased stool levels of CST. This might contribute to the reduced mucus layer in inflamed colon regions of IBD patients. CST feces levels might be a therapeutic target or biomarker for IBD.

## Acknowledgment

We thank thank Zbigniew Mikulski (La Jolla Institute for Immunology) and Kiek Verrijp (Radboud University Medical Center) for technical support with the histological stainings. We thank Roos Berbers (University Medical Center Utrecht) for supplying feces of healthy donors.

---

**Fig. S1. Calprotectin levels correlates with IBD activity**. **(A)** Calprotectin levels detected in feces of healthy donors (black) and Crohn’s disease (CD) patients in remission (grey) or flare-up (white). **(B)** Same as panel A, but now for calprotectin levels in feces of ulcerative colitis (UC) patients. Kruskal-Wallis *: P<0.05; **: P<0.01; ***P<0.001.

**Fig. S2. HE stain colon Crohn’s disease patient.** Representative images of hematoxylin and eosin (HE) stained colon cross-sections of non-inflamed **(A)** and inflamed **(B)** colon tissue of Crohn’s disease (CD) patients. Note the increased immune cell infiltration (blue cells) in the inflamed colon region.

**Table S1. Patient characteristics for feces measurements.** Characteristics of the patients at time of inclusion for stool measurements. As an indication for the extent of disease, the location has been scored according to the Montreal classification. For CD; L1 is ileal disease, L2 is colonic disease, L3 is ileocolonic disease, and L4 is isolated upper gastro-intestinal tract. For UC; E1 is distal to the rectosigmoid junction (proctitis), E2 is distal to the splenic flexure (left-sided) and E3 is proximal to the splenic flexure (extensive). Medication entails therapeutic categories in use for IBD; other therapies are not included. Some patients use more than one drug which all have been scored separately. Biologicals comprised only anti-TNF-α compounds. CD: Crohn’s disease, IBD: inflammatory bowel disease, UC: ulcerative colitis.

**Table S2. Patient characteristics for electron microscopy.** Characteristics of the patients at time of inclusion for electron microscopy on colonic biopsies. Indication for colonoscopy for the healthy control was a positive iFOBT. For patient for paired biopsies were taken, a and b.

## Data and materials availability

All data is available in the main text or the supplementary materials. Raw data is available upon request via Geert van den Bogaart and Sushil Mahata.

## Author contributions

E.M.M., S.K.M. and G.v.d.B designed the study and experiments. E.M.M, L.L, S.K.M., K.T. designed, analyzed and performed the experiments. L.L. and B.O. provided donor material. M.K. and J.F. performed the human colon electron microscopy. E.M.M. and G.v.d.B wrote the manuscript and all authors participated in discussing and editing of the manuscript.

